# Nuclear Speckles are Regulatory Hubs for Viral and Host mRNA Expression During HSV-1 Infection

**DOI:** 10.1101/2025.09.11.675548

**Authors:** Shani Nadav-Eliyahu, Chaya Bohrer, Alon Boocholez, Noa Kinor, Vesa Aho, Jennifer I.C. Benichou, Salla Mattola, Sami Salminen, Henri Niskanen, Minna U Kaikkonen, Maija Vihinen-Ranta, Yaron Shav-Tal

## Abstract

Herpes simplex virus type 1 (HSV-1) infection remodels the host nucleus, marginalizing chromatin and forming viral replication compartments (VRCs). Nuclear speckles, nuclear bodies enriched in RNA-processing factors, reposition around VRCs and undergo structural changes. While viral mRNAs are transcribed in VRCs and host transcription is largely suppressed, the nuclear routes used by viral and upregulated host transcripts and their relationship with nuclear bodies, remain unclear. We show that immediate-early (IE) viral transcripts uniquely accumulate in nuclear speckles prior to export, unlike early or late transcripts, revealing a selective nuclear speckle-dependent pathway. Similarly, host mRNAs upregulated during infection traffic into nuclear speckles after transcription. Moreover, nuclear speckles are structurally remodeled, marked by lncRNA MALAT1 removal and increased dynamics of the nuclear speckle core protein SRRM2. Lastly, we found that blocking mRNA export causes IE transcripts to accumulate in nuclear speckles, and that nuclear speckle disassembly severely impairs IE mRNA export, preventing downstream viral gene expression. These findings establish nuclear speckles as dynamic regulatory hubs that selectively facilitate the processing and export of IE viral mRNAs during HSV-1 infection.

**Significance statement:** This study reveals how herpes simplex virus type 1 (HSV-1) manipulates structures in the nucleus termed nuclear speckles, which are essential for processing of mRNA. We discovered that early viral messages specifically pass through these nuclear speckles before exporting out of the nucleus. We find that disassembling nuclear speckles severely limits viral RNA export. Moreover, certain host cell messages also rely on nuclear speckles during infection, suggesting a shared nuclear pathway for host and viral mRNAs during infection.

## Introduction

Herpes simplex virus type 1 (HSV-1) infection causes significant major alterations in nuclear architecture and disrupts numerous cellular processes. Upon entry of the viral genome into the nucleus, dramatic structural changes take place, including disassembly of the nucleolus (1) and some nuclear bodies (2–8), chromatin condensation, and disruption of the nuclear lamina (9). These changes coincide with the formation of viral replication compartments (VRCs), specialized nuclear regions where viral DNA replication occurs (10, 11). Initially, small VRCs emerge, which later expand. Live-cell imaging studies suggested that VRCs may merge from smaller VRCs (12, 13), however, a more recent study, using dual color FISH staining of the viral DNA, revealed that each VRC originates from a single incoming viral genome and that the VRCs territories are very stable (14). In conjunction, host chromatin is displaced toward the nuclear periphery (7, 15–17). These structural rearrangements also lead to transcriptional repression of many host genes, although some genes retain or even enhance their expression during infection (18–23). Recently, the importance of the spatial subnuclear localization of HSV-1 viral transcription in early infection times was discovered. Viral genomes tethered to the nuclear periphery upon entry, compared with those freely infecting the nucleus, were subjected to stronger transcriptional silencing. However, the subnuclear localization was found to be critical only during early infection (24).

In addition to chromatin displacement, VRC formation affects nuclear bodies such as nuclear speckles (25–28). These dynamic structures, enriched with splicing factors and other RNA-processing components, are redistributed during infection (6, 12) Importantly, HSV-1 VRCs have been shown to move and coalesce at nuclear speckles, suggestive of creating an optimized environment for viral RNA processing and export (12). This interaction underscores the functional reorganization of the host nucleus to facilitate viral replication while suppressing host antiviral defenses.

The functional role of nuclear speckles remains a topic of debate. Initially thought to be sites of pre-mRNA splicing, they are now recognized as hubs linked to high gene expression activity. Nuclear speckles may serve as reservoirs for splicing factors regulating their release near active genes or act as transit points for mRNAs to acquire additional processing and RNA-binding proteins (RBPs) before nuclear export (29–33). Recent studies have revealed the broader role of nuclear speckles in supporting transcriptional activity, particularly for genes near these structures. Genome-wide analyses have shown a strong correlation between gene expression levels and the distance of genes from nuclear speckles, with closer proximity corresponding to higher transcriptional activity (34–36). According to these studies, approximately one-fifth of the genome is situated near nuclear speckles, emphasizing their widespread influence on gene regulation (34). Moreover, organization of highly transcribed RNA Pol II transcribed genes at the periphery of nuclear speckles increases their splicing efficiency (36, 37). Disruption of nuclear speckles affects the expression of thousands of genes, emphasizing their importance in maintaining cellular homeostasis (38).

It has been suggested that HSV-1 exploits nuclear speckles to optimize its replication cycle. The proximity of VRCs to nuclear speckles assumably enhances the recruitment of RNA-processing factors and facilitates the export of viral transcripts (12). For instance, SRSF3 and SRSF7 splicing factors, which are nuclear speckle components, were found to be important for viral RNA export, with their knockdown significantly impairing viral yields and leading to nuclear retention of polyadenylated viral RNA (39). HSV-1 infection leads to the redistribution of other nuclear bodies. For example, infection conditions affect the association of SRSF2 with various paraspeckle factors, thereby influencing histone modifications near viral genes and the splicing of the long-noncoding RNA NEAT1 (5, 40). Interestingly, KSHV induces enlarged, structurally distinct paraspeckles near viral replication centers, which likely serve as RNA processing hubs essential for lytic replication (41, 42). Nuclear speckles and paraspeckles are vital also in other viruses, such as HIV and influenza, which leverage these nuclear bodies to facilitate transcription and splicing of their RNA (43, 44). The influenza NS1 protein was found to promote M viral mRNA post-transcriptional splicing and export via nuclear speckles (43). The host nuclear pore component, TPR, and the TREX2 complex are essential for influenza mRNA nuclear export (45). HIV VRCs traffic to and accumulate within nuclear speckles. Interestingly, HIV-1 shows a strong preference for integrating into genomic domains associated with nuclear speckles (44).

Despite extensive evidence of nuclear reorganization during viral infection, the precise nuclear localization and trafficking pathways of viral and host cellular mRNAs following transcription remain poorly understood. In this study, we aimed to visualize endogenous viral and host mRNAs within the nucleus during HSV-1 infection and characterize their interactions with nuclear speckles. By employing RNA imaging techniques in fixed and living cells, we demonstrate that viral transcription occurs exclusively within viral replication compartments and not within nuclear speckles. Importantly, immediate-early (IE) viral transcripts uniquely accumulate within nuclear speckles following transcription, whereas early (E) and late (L) transcripts do not display this localization pattern. We also discovered that certain host genes continue to be actively transcribed near nuclear speckles during infection, with their mature transcripts subsequently accumulating in these nuclear structures. Furthermore, significant compositional and dynamic changes occur in nuclear speckles during infection, including the depletion of the long non-coding RNA MALAT1 and increased mobility of the core speckle protein SRRM2. Finally, we establish that nuclear speckles function as critical intermediate hubs for IE viral mRNAs prior to export, highlighting their essential role in viral RNA processing and nuclear export pathways.

## Materials and methods

### Cell Culture

U2OS human osteosarcoma cells were maintained in low-glucose Dulbecco’s modified Eagle’s medium (DMEM) (Biological Industries, Beit-Haemek, Israel) containing 10% FBS (HyClone Laboratories, Logan, UT, USA). Cells were grown at 37°C and 5% CO_2_. HFF-1 cells (human foreskin fibroblasts; provided by Ron Goldstein, BIU, Ramat Gan, Israel) were maintained in high-glucose DMEM (Biological Industries) containing 10% FBS. Mycoplasma tests were carried out periodically on all cell lines. For each experiment, cells were stained with trypan blue solution (Sigma T8154), counted, and seeded on coverslips. For transcription inhibition, Flavopiridol (Sigma F3055) was added to the medium to a final concentration of 1 nM at the last hour of infection.

### Knock-in cell lines

For the knock-in of the mNeongreen fluorescent protein into the C-terminus of the endogenous *NPM1* (B23) gene, we used the Mammalian PCR-tagging protocol for tagging the C-termini of genes (46). The pMaCTag-P07+MS2v5 plasmid containing the mNeongreen coding region and 24xMS2v5 sequence was used. For knock-in of the mScarlet fluorescent protein into the C-terminus of the endogenous SRRM2 gene, a pMaCTag-Z12 (#120055) plasmid was used. The online oligo design tool www.pcr-tagging.com (46) was used for searching PAM sites and for designing the forward (M1) and reverse (M2) tagging oligos specific for impLbCas12a (Addgene #137715). M1 and M2 oligos, targeting NPM1 and SRRM2, were obtained from IDT.

NPM1 M1:

ATGTCTATGAAGTGTTGTGGTTCCTTAACCACATTTCTTTTTTTTTTTTTCCAGGCTATTCAAGATCTCTGGCAGTGGAGGAAGTCTCTTTCAGGTGGAGGAGGTAGTG

NPM1 M2:

TTACAGAAATGAAATAAGACGGAAAATTTTTTAACAAATTGTTTAAACTATTTTCAAAAAAGTTTAAACTATTTTCTTAAAATCTACACTTAGTAGAAATTAGCTAGCTGCATCGGTACC

SRRM2 M1:

CGGCCCCATTTTGGGAGTGGCCCAGAAACTGGCCTTGAGGGCTGGGGTGGGAACTCCCTGTTGACCCATATCTTCTCTTGCAGGTCTCCATCAGGTGGAGGAGGTAGTG

SRRM2 M2:

GGACACTCCTTGTGGCTCCAGAGCATTGGGTGTGGTGGAATCCCCCAAAGACAATAAAAAACAATTTATGGAGACCTGCAAATCTACACTTAGTAGAAATTAGCTAGCTGCATCGGTACC

The cassette was amplified with M1 and M2 using pMaCTag-P07-NG-MS2v5/ pMaCTag-Z12 as a template. To generate U2OS Tet-On cells expressing endogenously tagged B23-mNeongreen/SRRM2-mScarlet cells, ∼10^6^ cells were seeded 24 hrs before transfection. Plasmid impLbCas12a and the mNeongreen-NPM1/ mScarlet-SRRM2 PCR cassette were electroporated (Bio-Rad) into the cells. 72 hrs later, the cells were trypsinized and Puromycin/Zeocin selection was added. After 7–14 days, cells were examined for mNeongreen/mScarlet fluorescence and seeded into a 96-well plate for the generation of single clones. After another 1–2 weeks, positive clones were transferred to a 24-well plate for further analysis. Genomic DNA from U2OS Tet-On and the knock-in B23-mNeonGreen/SRRM2-mScarlet cells were extracted with the Wizard® Genomic DNA Purification Kit (Promega). Clones were genotyped by PCR for homozygous insertion of tags, and the PCR product was sent for sequencing verification. For PCR verification of the SRRM2 knock-in primers used were: forward ATGAGACACCGCTCCTCCAGGT (on the *SRRM2* gene), reverse GAGTTTGGACAAACCACAACTAGAATGC (on the cassette). For PCR verification of the *NPM1* (B23) knock-in, primers used were: Forward CTAGAGTTAACTCTCTGGTGG (on the *NPM1* gene), reverse GGCCCTCGTTTGACGTAT (on the cassette).

### Viral infection

U2OS cells were grown on coverslips for 24 hrs and were then infected with the HSV-1 wild type (WT, 17^+^), ICP4-YFP (PMC369473), and VP26-mCherry virus (17^+^ Lox-CheVP26) (47–49) using a multiplicity of infection of 5 in FBS-free DMEM culture medium. HFF cells were infected with the ICP4-EYFP at a multiplicity of infection of 10. At 1 hr post-incubation with either virus, the culture medium was replaced with the same culture medium containing 10% FBS to remove unabsorbed virus. Then cells were incubated for 1-8 hrs according to the experiment. For live-cell imaging and FRAP, DMEM was replaced with phenol-free L-15 (Gibco 21083027) containing 10% FBS for the entire imaging period. After the required time, cells were fixed with 4% PFA.

### Plasmids and transfections

For generating the A1-Halo-MCP plasmid, we first amplified the Halo sequence from phage-nls-ha-tdMCP2-halo (Gal Haimovich, Weizmann Institute) with the addition of homolog ends targeting the vector, by PCR using HiFi HotStart ReadyMix (Roche KK2602). Then, we amplified the previously described YFP-A1-MCP plasmid (50) to a linear vector, excluding the YFP sequence, by PCR using HiFi HotStart ReadyMix (Roche KK2602). The Halo insert and A1-MCP linear vector were then annealed together using Gibson assembly LigON Master Mix (Biotech applications E1050) to get the final A1-Halo-MCP plasmid.

Primers used for amplifying the Halo sequence:

Halo Fwd:

GGGAGGAAATTTTGGAGGCAGAAGCTCTGGACCCTATGGCGCAGAAATCGGTACTGGCT

Halo Rev:

AAGCCATTAAGCCTGCTTTTTTGTACAAACTTGTTTGATAACCGGAAATCTCCAGAGTAGACAG

Primers used for amplifying the A1-MCP linear vector:

A1-MCP Fwd: TATCAAACAAGTTTGTACAAAAAAGCAGGCT
A1-MCP Rev: GCCATAGGGTCCAGAGCTT

For generating the pMaCTag-P07-NG-MS2v5 plasmid, we amplified the 24xMS2v5 sequence from the MSV5 vector (#84561) and added SpeI restriction sites at both ends using PCR with KAPA HiFi HotStart ReadyMix (Roche KK2602). The amplified sequence was then subcloned into the PMacTgp07 plasmid (#124788). Primers used for amplifying 24xMS2 with SpeI restriction sites:

MS2v5 Fwd: TATACTAGTGGTAACCTACAAACGGGTGGA
MS2v5 Rev: TATACTAGTGATATGAGATCTGAGGTGTTTGATGT.

Cerulean-Dbp5-DN (51), TNPO3-cargo binding domain and cerulean-Clk1 (30) were previously described.

For transfection, the ViaFect reagent (Promega E4981) was used according to manufacturer’s protocol. Cells were transfected with the plasmid 24 hrs prior to infection. Transfection of A1-Halo-A1-MCP in U2OS cells for live-cell imaging was done by electroporation 24 hrs prior to infection.

### Single molecule RNA FISH

Stellaris protocol: Cells were seeded on 13 mm coverslips, infected and fixed for 20 min in 4% PFA, then left overnight with 70% ethanol at 4°C. Coverslips were then washed twice with 10% formamide (Thermo-Fisher 327235000). Probes (RL2 Exons/RS1/UL30) were diluted in 10% formamide and 10% Saline-sodium citrate (SSC) hybridization buffer and hybridized with cells overnight at 37°C in a dark chamber. The next day, cells were washed twice with 10% formamide for 15 min at 37°C and then washed with 1x PBS. To reduce photobleaching, the cells were submerged in glucose oxidase (GLOX) buffer (pH = 8, 10mM, 2x SSC, 0.4% glucose), supplemented with 3.7ng of glucose oxidase (Sigma G2133-10KU) and 1µl Catalase (Sigma 3515) prior to imaging (18806792, 12791999).

FLAP FISH: We implemented the FLAP-FISH protocol as published (52). Three sets of FLAP tails were used: X (CACTGAGTCCAGCTCGAAACTTAGGAGG), Y (AATGCATGTCGACGAGGTCCGAGTGTAA), and Z (CTTATAGGGCATGGATGCTAGAAGCTGG). X-tails were conjugated with either Cy3 (Ex 548/Em 566) or Cy5 (Ex 647/Em 670) fluorophores. Y and Z tails were conjugated with Cy5 fluorophores. 28 nt tails were added to the 5’-end of each probe designed, the reverse-compliment sequence of the fluorescent tail. After diluting the probes to 100 µM, they were mixed and then diluted to a concentration of 0.833 µM. Next, the protocol described in (52) was used to hybridize the probes with their fluorescent tails. FLAP experiments were performed according to the Stellaris adherent cell protocol. Probe sets used: RL2 intron 1 (Y), RS1 (Y), US1 exons (Y), US1 intron 1 (X), NXF1 exons (X), NXF1 introns 1-2-3 (Z), SRSF2 (Y), SRSF7 (Z), MALAT1 (X).

### Immunofluorescence

Primary antibodies were added to the hybridization buffer together with probes at the suitable dilution: Rabbit anti-SRRM2 [Abcam; ab122719]: 1:400, mouse anti-ICP4 [ab6514]: 1:1000, mouse anti-NXF1 [ab129160]: 1:250, mouse anti-Pol II Ser-5 (H14), mouse anti-Pol II Ser-2 (H5) hybridomas (36). Secondary antibodies: goat anti rabbit IgG H&L Alexa fluor 488 [ab150077], donkey anti-rabbit IgG H&L Alexa flour 647, anti IgM-Cy3 [Jackson ImmunoResearch, West Grove, PA), donkey anti-mouse IgG H&L Alexa fluor 750 [ab175738] were applied to cells at the end of the FISH procedure diluted 1:1000 in 1xPBS. Nuclei were counterstained with Hoechst (33342).

### Fluorescence Microscopy and Image Analysis

Wide-field fluorescence images were obtained using the CellSens system based on an Olympus IX81 fully motorized inverted microscope (60x Planpon O objective 1.42 [NA]) fitted with an Orca-Flash4.0 #2 camera (Hamamatsu, Bridgewater, NJ, USA), rapid wavelength switching, and driven by the CellSens v3.2 software (Olympus, Tokyo, Japan); or the CellR system based on an Olympus IX81 fully motorized inverted microscope (60x PlanApo objective 1.42 [NA] Olympus) fitted with an Orca-AG CCD camera (Hamamatsu), rapid wavelength switching, and driven by the CellR v1.2 software (Olympus). For time-lapse imaging, cells were plated on glass-bottomed tissue culture plates (MatTek, Ashland, MA, USA) in DMEM medium containing 10% fetal calf serum at 37°C. The microscope is equipped with an on-scope incubator which includes temperature and CO_2_ control (Life Imaging Services, Reinach, Switzerland). For long-term imaging, several cell positions were chosen and recorded by a motorized stage (Scan IM; Märzhäuser, Wetzlar-Steindorf, Germany). Cells were typically imaged in three dimensions (3D) (11 Z planes per time point, step size = 1µm) every 5 min or 10 min (depending on the experiment). Live imaging of NPM1 transcripts was imaged in single Z plane every 400 ms. For nuclear speckles size and volume quantification, mock and 6 hpi infected cells were imaged in 3D (37 Z planes, step size = 0.3 µm).

Live-cell imaging and 3D stacks were deconvolved using Huygens Essential v18.10 software (Scientific Volume Imaging, Hilversum, Netherlands). Quantification of nuclear speckles size and volumes was performed using the Imaris softwere. Colocalization analysis of two channels was performed using an ImageJ Macro (Shav-Tal lab, Ramat Gan, Israel).

To analyze the localization of mRNA foci, nuclear speckles and the replication compartment, the replication compartment was first segmented using either Otsu’s or maximum entropy algorithm for the ICP4 channel (53, 54). Otsu’s algorithm was used as a default, and if it did not produce a satisfactory result, judged by visual inspection, the maximum entropy algorithm was used instead. The nuclear speckles were segmented by setting the threshold of the SRRM2 channel to twice its mean value in the nucleus. For the mRNA channel, the threshold was manually adjusted for each cell. FRAP experiments were performed using a Leica Laser scanner based on the Leica Total Internal Reflection Fluorescence (TIRF) microscope (63x HC PL APO objective 1.40 [NA]) with the LASX v3.7.5 software (Leica,), as previously described (30). Before bleaching, cells were imaged in the YFP channel for the detection of ICP4-EYFP. SRRM2 signal at a specific nuclear speckle was photobleached using the 553 nm laser. 5 pre-bleach images were acquired. 200 post-bleach images were acquired every 0.5 sec. For each time-point, the background taken from a ROI outside of the cell was subtracted from all other measurements. T(t) and I(t) were measured for each time-point as the average intensity of the nucleus and the average intensity in the bleached ROI, respectively. The average of the pre-bleach images used as the initial conditions are referred to as Ti = nuclear intensity and Ii = intensity in ROI before bleaching. Ic(t) is the corrected intensity of the bleached ROI at time t (15205532,10766243):

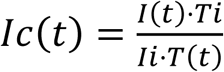

Data from at least 3 experiments and at least 30 nuclear speckles for each treatment were collected and the averaged FRAP measurements were fitted to a curve.

### Western blotting

Cells were infected for the desired time and washed in cold PBS, and proteins were extracted in pierce IP lysis buffer (Thermo 87787) containing 10 mM Na-flouride, 1 mM Na-orthovanadate, protease inhibitor cocktail (Sigma) and 1 mM PMSF, and placed on ice for 20 min. The resulting lysate was centrifuged at 10,000 rpm for 10 min at 4°C. 30 μg/μl of protein/lane was run on SDS-polyacrylamide gels and transferred to a nitrocellulose membrane (0.45 μm). The membrane was blocked in 5% BSA, and then probed with mouse anti-NXF1 primary antibody [ab129160] for 2 hrs at room temperature (RT), followed by incubation with HRP-conjugated goat anti-rabbit IgG (Jac-111-035-144, Jackson) for 1 hr at RT. For loading control, the blots were reblotted with an anti-tubulin antibody (DSHB), followed by incubation with HRP-conjugated goat anti-mouse IgG (115-035-062, Jackson) for 1 hr at RT. Immunoreactive bands were detected by the Enhanced Chemiluminescence kit (ECL Thermo 34580).

### RNA extraction and Real-Time PCR

Cells were infected with the HSV-1 (MOI =5) for 6 hrs or infected for 6 hrs and treated with Flavopiridol at the last hour of infection. Total RNA was extracted from cells using TRI Reagent following Direct-zol RNA MiniPrep (ZYMO research).

After reverse transcription using qScript cDNA Synthesis Kit (Quanta Biosciences, MA, USA), the cDNA was amplified using the following primer pairs:

UL30: Forward primer: 5’- CGAGTTTAACTGTACGGCGG- 3’
UL30 Reverse primer: 5’- CCCGCCTTGCATTCGATATC- 3’
UL36 Forward primer: 5’- GTCCTCGCCTTTCTCATCGA-3’
UL36 Reverse primer: 5’- TCAGGGTAAACTCCAGCAGG-3’
ICP4 Forward primer: 5’- AGGTGACCTACCGTGCTAC-3’
ICP4 Reverse primer: 5’- CTTGTTCTCCGACGCCATC
18S Forward primer: 5’- CATGAACGAGGAATTCCCAGTAA- 3’
18S Reverse primer: 5’- GATCCGAGGGCCTCACTAAAC- 3’
Homo sapiens ubiquitin conjugating enzyme E2 I (UBE2I)
Forward primer: 5’- GTGTGCCTGTCCATCTTAGAG- 3’
Reverse primer: 5’- GCTGGGTCTTGGATATTTGGTTC- 3’

Real-time RT-PCR was performed using PerfeCTa® SYBR® Green FastMix®, ROX™ (Quanta Bio Sciences, MA, USA) on a CFX-96 system (Bio-Rad, CA, USA). Analysis was performed with the Bio-Rad CFX manager. Relative levels of mRNA expression were measured as the ratio of the comparative threshold cycle (CT) to internal controls (UBEL2I and 18S) RNA.

### Statistical analysis

All statistical analyses were performed using R statistical software v4.1 (Oxford Protein Informatics Group, Oxford, United Kingdom) (R Core Team (2021). R: A language and environment for statistical computing. R Foundation for Statistical Computing, Vienna, Austria. https://www.R-project.org/), or by Prism GraphPad software.

For analysis of the FRAP data, three-parameter asymptotic regression was used. Each nuclear speckle recovery data was fit to a curve defined by the equation where X is time and Y is the relative intensity, a (plateau) is the maximum attainable intensity, b (init) is the initial Y value (at time=0) and c (m) is proportional to the relative rate of increase for intensity when time increases. Each parameter (init, m and plateau) was then compared between mock, 2 hr and 6 hr with a one-way nested ANOVA, followed by pairwise comparisons of mean values between treatments. Finally, *p*-values were adjusted for multiple comparisons with the Benjamini-Hochberg (FDR) procedure. Outliers were removed using the Interquartile-range criterion.

A one-way ANOVA was performed to analyze the difference between all 3 viral mRNAs (RL2, RS1, UL30) overlapping with nuclear speckles, the effect of treatment (Mock, 2 hpi, 4 hpi and 6 hpi) on the percentage of cells with dispersed MALAT1, and the levels of NXF1 protein in mock, 2, 4, and 6 hpi detected by western blotting, followed by Tukey’s post hoc analysis. For analysis of the viral mRNAs overlap with nuclear speckles at early and late stages, a two-tailed *t*-test was performed. A one-way ANOVA was performed to analyze the difference between the decrease in mRNA levels after falvopiridol treatment (ΔΔCt values) of each viral mRNA measured by qPCR. A two-way ANOVA was performed to analyze the effect of treatment (control and Dbp5) and transcripts (RL2, RS1 and UL30) on viral mRNA foci counts, followed by Tukey’s post hoc analysis. To assess group differences in number of NS and volume, one- and two-way mixed-design ANOVA tests were conducted using linear mixed-effects models. The models included group (mock and 6 hpi) as a fixed effect and rep as a random effect to account for the dependency between cells from same biological replicates. Fixed effect significance was tested using Type III ANOVA, and pairwise post hoc comparisons between groups were conducted using estimated marginal means with Tukey adjustment for multiple comparisons.

### Global run-on sequencing (GRO-seq)

Global run-on sequencing (GRO-seq) analysis of viral and cellular transcripts was performed in HeLa cells (ATCC® CCL-2) at 4 and 8 hpi, as previously described (55).

## Results

### Immediate-early viral mRNAs localize in nuclear speckles

Nuclear organization is extensively modified upon HSV-1 infection. Nuclear speckles redistribute and coalesce at nuclear sites associated with viral genome replication. However, the spatial distribution of viral and cellular transcripts within the nucleus, as well as the role of nuclear speckles in gene expression during HSV-1 infection, remains poorly understood. To investigate this, we first established the detection of viral mRNAs in infected cells using RNA FISH. Probe sets were designed to target different viral mRNAs: RL2, encoding the ICP0 protein; RS1, encoding the ICP4 protein; and UL30, encoding the catalytic subunit of viral DNA polymerase. *RL2* and *RS1* are classified as immediate early (IE) genes, while *UL30* is an early (E) gene. Notably, *RL2* (ICP0) contains introns, and therefore, these transcripts undergo splicing, which is considered unique, as most viral genes lack introns. The 3 viral transcripts were detected in human U2OS cells infected for 1-6 hrs (Figure 1A, Supplementary Figure 1A). No RNA labeling was observed in mock-infected cells, whereas RNA labeling and increasing levels of viral mRNA were evident in cells infected over 1-6 hrs. At 1 hr post-infection (hpi), abundant RL2 and RS1 mRNAs were observed in the cytoplasm, along with bright foci in the nucleus (Supplementary Figure 1A), likely corresponding to active sites of viral mRNA transcription. UL30 mRNA became detectable at 3 hpi and continued to increase in intensity over time. As expected, by 4 hpi, viral mRNA was observed in both the cytoplasm and in larger nuclear regions, specifically within the viral replication compartments (VRCs), as identified by Hoechst staining (Figure 1A, Supplementary Figure 1A). Quantification of the viral mRNA demonstrates increased mRNA levels in the VRCs, as VRCS size are larger (Supplementary Figure 1B). Notably, as infection proceeded (6 hpi), cells with large VRCs, indicating a more advanced infection stage, showed distinct bright foci containing RL2 and RS1 mRNAs that surrounded the VRCs. This phenomenon was observed exclusively for RL2 and RS1, while UL30 mRNA remained primarily in the cytoplasm and within the VRCs in the nucleus (Figure 1A, Supplementary Figure 1A). To determine whether the phenomenon of viral mRNA foci formation is specific to IE genes, we infected the cells for 7 and 8 hrs and labeled for representative viral mRNAs from each gene class: immediate-early - *RL2*; early - *UL30*; and late - *UL36*. Viral mRNA foci were observed exclusively for the IE mRNAs of RL2, but not for UL30 or UL36, even at longer infection times (Supplementary Figure 1C).

**Figure 1:**
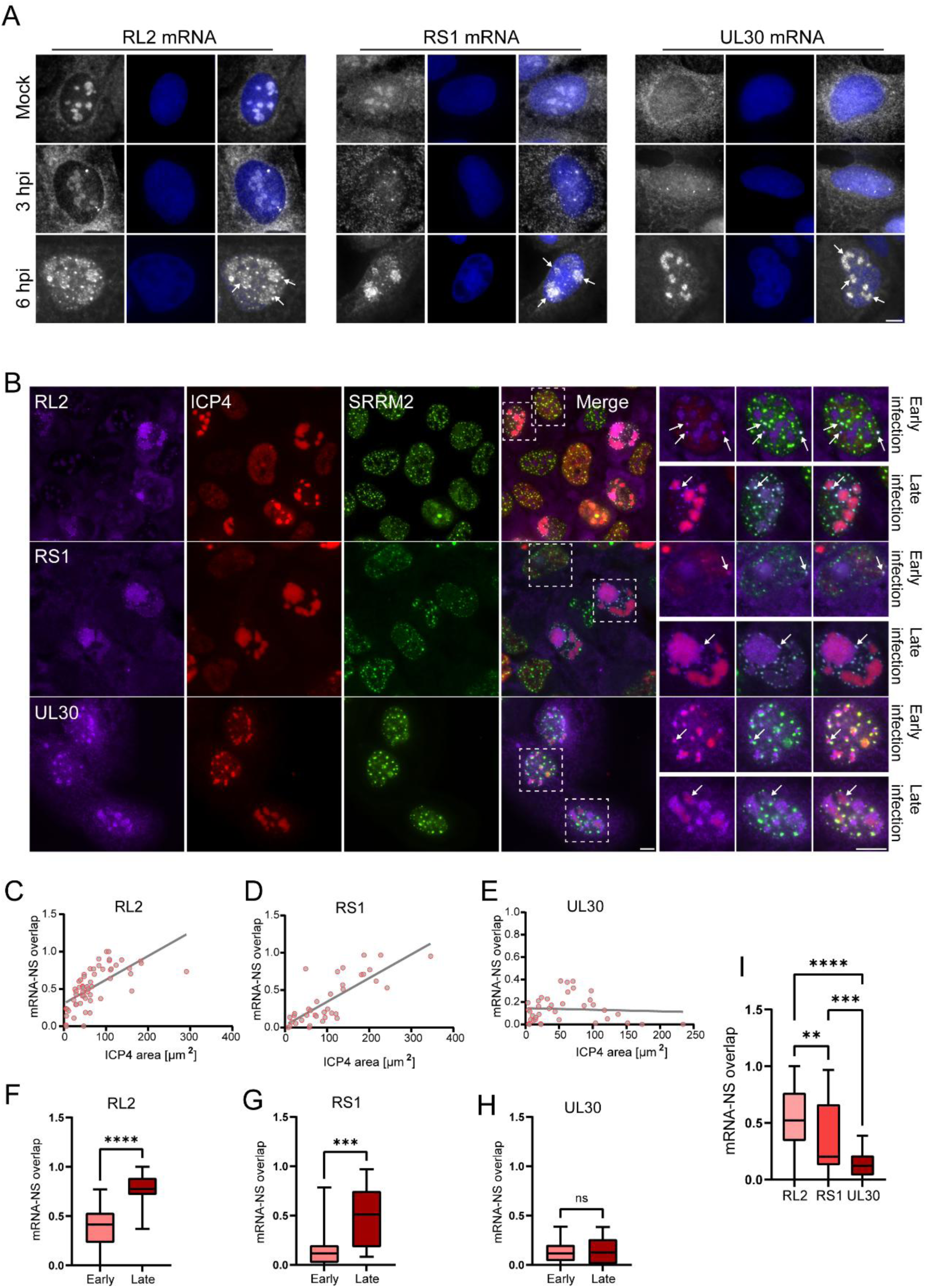
Viral mRNAs are transcribed near nuclear speckles and IE *RL2* and *RS1* viral mRNA accumulate within. (A) U2OS cells were mock infected and infected for 3 and 6 hrs (MOI=5) and labeled with fluorescent probes to RL2/RS1/UL30 viral mRNAs by RNA FISH (gray). Viral mRNAs are detected in the nucleus and cytoplasm and at 6 hpi also in bright foci inside the nucleus (in mock infected cells there is background signal of non-specific binding of probes within nucleoli). The nucleus was counterstained with Hoechst (blue). The entire field of the cells shown are presented in Supplementary figure 1. (B) U2OS cells were infected for 6 hrs (MOI=5) and labeled with fluorescent probes for the viral mRNAs RL2/RS1/UL30 (purple). ICP4 protein marks the VRCs (red), and SRRM2 marks nuclear speckles (green). Enlarged images show cells at an early infection stage where viral mRNAs are found near nuclear speckles and at a late infection stage where IE viral mRNAs are found localized within nuclear speckles (infection stage determined by VRC volume-ICP4 area). Scale bars= 10 µm. (C-E) Plots depicting the overlap of nuclear speckles (NS) with viral mRNA (C) RL2, (D) RS1 and (E) UL30 (n=59, 37, 38 cells, respectively). (F-H) Nuclear speckles overlap with viral mRNA plots are divided to early and late infection stages; (F) RL2, (G) RS1, (H) UL30. Nuclear speckles overlap with RL2 (*P* < 0.0001) and RS1 (*P* = 0.001) viral mRNAs are statistically different between early and late infection, while there is no significant difference with UL30 (*P* = 0.8198). Error bars show the standard error of the mean. (I) Nuclear speckles overlap with all three viral mRNA and show a statistical difference between all 3 transcripts: RL2-RS1 *P* = 0.0046, RL2-UL30 *P* < 0.0001, RS1-UL30 *P* = 0.0004.

Nuclear speckles are known to undergo reorganization in response to HSV-1 infection, relocating to the nuclear periphery and particularly around viral replication compartments (VRCs) (6, 12). Live-cell imaging of cells in which the endogenous nuclear speckle core protein SRRM2 was tagged with mScarlet, and infected with HSV-1 ICP4-EYFP, demonstrated the reorganization of the nuclear speckles around VRCs in real time (Movie S1). The accumulation of RL2 and RS1 mRNAs during 6 hpi closely mirrored the distribution pattern of nuclear speckles. At this time, the colocalization of RL2 and RS1 mRNAs with the nuclear speckle marker SRRM2 was observed (Figure 1B, Supplementary Figure 2), and cells at different stages of infection were evident, as indicated by ICP4 staining, which marks the VRCs. Interestingly, in early-stage infected cells, several nuclear RNA foci were observed co-localized with ICP4-positive early VRCs. These foci were also found associated with nuclear speckles, namely, they were closely positioned to them. At more advanced infection stages, when VRCs had expanded, viral mRNAs were detected not only within the VRCs but also in foci surrounding them, where they co-localized with nuclear speckles but did not overlap with ICP4 (Figure 1B, Supplementary Figure 2). These observations suggest that early in infection, small viral sites of transcription co-localize with ICP4 (closely positioned near nuclear speckles), and as infection progresses, these sites increase in size, forming within the larger VRCs that eventually occupy most of the nuclear space. However, in addition to the VRCs, viral mRNAs accumulated in nuclear speckles. This phenomenon was observed for the immediate early (IE) mRNAs RL2 and RS1 but not for the early (E) mRNA UL30, which did not co-localize with nuclear speckles and was confined to the VRCs (Figure 1B, Supplementary Figure 2). Measurements of the nuclear speckles overlapping with the viral mRNA demonstrated that overlap of the IE viral mRNAs (RL2 and RS1) increased as ICP4 regions became larger (Figure 1C,D) e.g. at late infection stages (Figure 1F,G). Notably, this overlap with nuclear speckles was unique for the IE viral mRNAs RL2 and RS1 and not UL30 (Figure 1E,H,I). However, there was also a significant difference in the overlap of nuclear speckles with either RL2 or RS1 mRNAs (Figure 1I). This can be explained by the fact that RL2 contains introns and requires splicing, whereas RS1 does not, leading to higher localization of RL2 mRNA to nuclear speckles. These findings were also observed in HFF cells (Supplementary Figure 3A). In conclusion, the immediate early (IE) viral mRNAs RL2 and RS1 initially localize in small VRCs and are later observed in both VRCs and nuclear speckles, whereas the viral early mRNA UL30 remains confined to VRCs.

To further investigate whether the viral mRNA foci co-localizing with nuclear speckles represent active transcription sites or areas of mRNA accumulation, perhaps as a route to the cytoplasm, we sought to specifically examine the RL2 viral mRNA and another immediate early (IE) mRNA, US1, which encodes the ICP22 protein. The RL2 pre-mRNA contains introns, and US1 mRNA was also identified as a viral transcript that contains introns and can potentially undergo alternative splicing (56). To analyze their transcriptional dynamics, we generated two probe sets for both RL2 and US1: one targeting the exonic regions of the transcript and another set detecting the intronic regions of the mRNA (Figure 2A,B). Thus, we were able to differentiate between pre-mRNA (unspliced) and mature mRNA (spliced). As expected, introns of both genes were detected exclusively in the nucleus, with no cytoplasmic signal (Figure 2A,B). IE US1 viral mRNA was also detected in VRCs and in foci co-localized with nuclear speckles at late infection stage, further supporting the hypothesis that viral mRNA association with nuclear speckles is unique to IE genes (Figure 2B). Notably, introns were localized to both the VRCs and to the foci associated with nuclear speckles (Figure 2A,B). This suggests that viral transcription occurs in the VRCs and that nuclear speckles may also be involved in the transcriptional or post-transcriptional processing of RL2 and US1, either as sites of active transcription or splicing.

**Figure 2:**
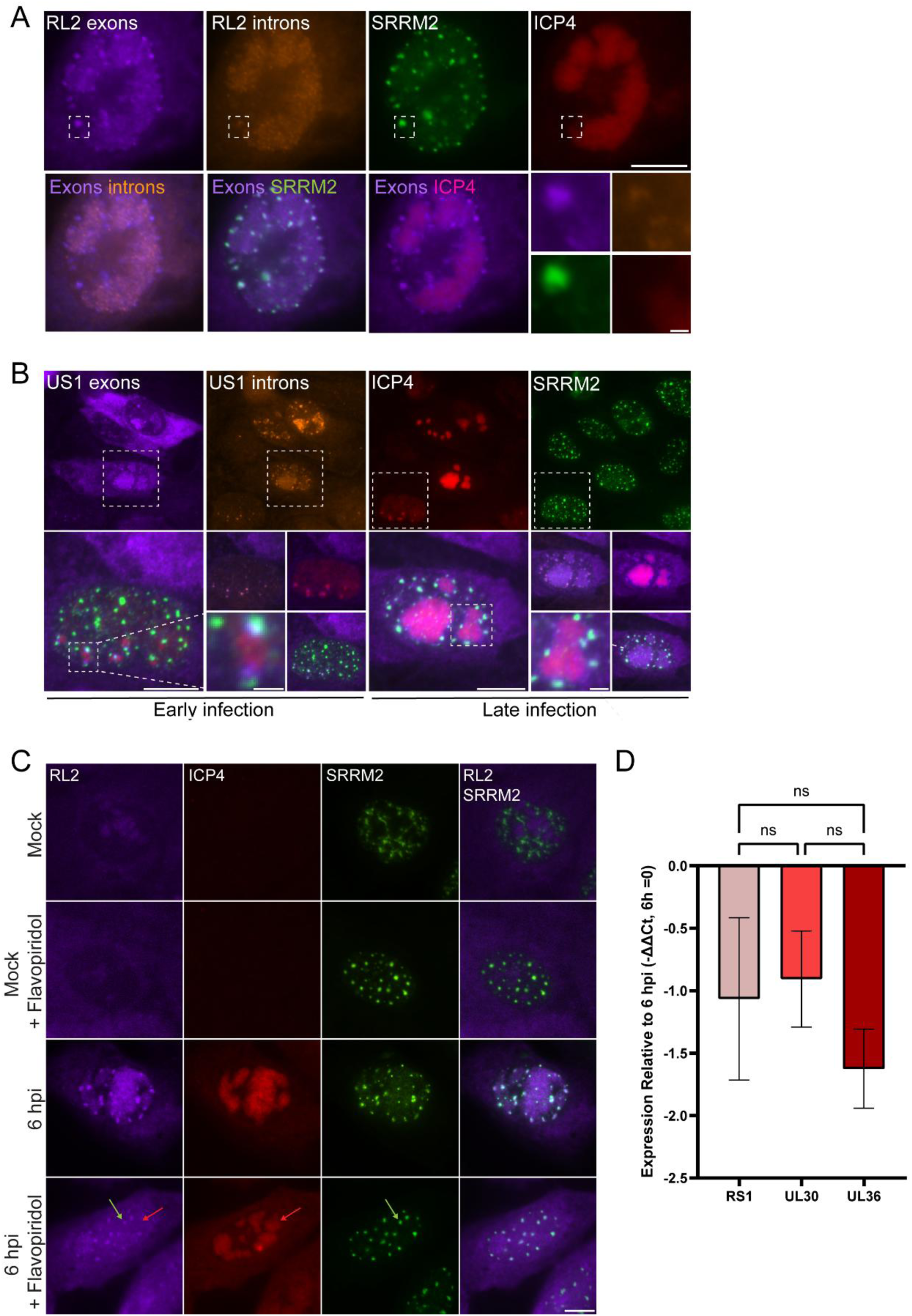
Nuclear speckles are sites of IE viral mRNA accumulation, not transcription. (A) U2OS cells were infected for 6 hrs (MOI=5) and then labeled with fluorescent probes for RL2 viral mRNA by RNA FISH, using specific probes to RL2 exons (purple) and introns (orange). SRRM2 marks nuclear speckles (green) and ICP4 marks VRCs (red). (B) U2OS cells were infected for 6 hrs (MOI=5) and labeled for US1 viral mRNA using specific probes to US1 exons (purple) and introns (orange). SRRM2 marks nuclear speckles (green) and ICP4 marks VRCs (red). Enlarged images show cells at an early infection stage and at a late infection stage (determined by VRC volume). (C) U2OS cells were infected for 6 hrs (MOI=5) and treated with Flavopiridol at the last hour of infection. Cells were labeled with fluorescent probes for RL2 viral mRNA (purple). SRRM2 marks nuclear speckles (green). ICP4 (red). Scale bar= 10 µm. (D) Plot of qRT-PCR demonstrating the change in the expression of three viral transcripts (RS1, UL30, UL36) under transcription inhibition treatment for 1 hr, relative to 6 hpi (ΔΔCt). All three viral transcripts demonstrate a decrease in their levels under transcription inhibition with no statistical difference between them (n=3).

To further clarify whether the viral mRNAs in nuclear speckles represent nascent mRNAs being transcribed within these foci or that the mRNAs are accumulating there post-transcriptionally, transcription was inhibited and the localization of the mRNAs was examined. At 3 hpi, the transcription inhibitor Flavopiridol was added during the final hour of infection. Inhibition led to rounding up of nuclear speckles as expected, and to a significant reduction in RL2 mRNA inside VRCs. However, bright RL2 mRNA foci colocalized with nuclear speckles remained visible (Figure 2C). These findings indicate that viral mRNA is primarily transcribed in the VRCs and subsequently accumulates in nuclear speckles. The presence of introns in these nuclear speckles-associated foci but not in the nucleoplasm suggests that the transcripts likely undergo post-transcriptional splicing within the nuclear speckles. To further clarify the localization of IE transcripts in nuclear speckles compared to the dis-localization of the E and L transcripts, we examined whether there might be a difference in the stability of the viral transcripts. RNA was extracted from 6 hpi infected cells and from 6 hpi infected cells treated with flavipiridol during the last hour of infection, and using qRT-PCR the levels of the viral mRNAs under the different treatments were quantified. No significant difference in the change of expression between three viral transcripts was found (Figure 2D), suggesting there is no variation in the transcript’s stability.

### Upregulated host mRNAs localize in nuclear speckles during HSV-1 infection

Discovering viral mRNA localization in nuclear speckles led us to examine whether host gene expression would also be associated with these nuclear bodies during HSV-1 infection. Most host cellular genes are suppressed due to viral host gene shut-off (57). To identify specific cellular genes upregulated during HSV-1 infection, we reanalyzed our previous GRO-seq data from HeLa cells (55). The analysis revealed that HSV-1 infection increased the mRNA levels of the transcription of nuclear export factor 1 (NXF1/TAP), SRSF2 and NPMP1 (Supplementary Figure 4A). The probe sets targeting the exonic and intronic regions of NXF1 mRNA, verified a significant increase in NXF1 mRNA levels at early-stage infection (indicated by spotted nuclear distribution of ICP4) compared to mock-treated cells (Figure 3A). In addition to single transcripts, larger nuclear foci of NXF1 mRNA signal, most likely sites of NXF1 transcription, were detected. Examination of the pre-mRNA (NXF1 introns) revealed that, as expected, unspliced NXF1 transcripts predominantly localized to these large NXF1 mRNA foci, as most splicing events occur co-transcriptionally (58). However, additional foci of NXF1 mRNA lacking introns were detected, indicating the assembly of mature, spliced transcripts (Figure 3A). Thus, by employing a two-color exons/introns labeling approach, we were able to identify and differentiate between NXF1 transcription sites and sites of accumulation of mature NXF1 mRNA. Notably, NXF1 foci were co-localized with nuclear speckles (Figure 3A). This association was observed both for intron-containing foci (transcription sites) and for intron-less foci (accumulation of mature mRNA) (Figure 3B). However, close examination of NXF1 intron labeling and SRRM2 localization reveals that while the transcription site of NXF1 is closely associated with nuclear speckles, it does not overlap with them. In contrast, exon labeling within the same foci appears larger and is more co-localized with nuclear speckles, likely due to its larger distribution (Figure 3A). This suggests that transcription occurs adjacent to nuclear speckles rather than within them. However, mature transcripts, whether still at the transcription site or after release, are found inside nuclear speckles. A similar pattern was observed in HFF-infected cells (Supplementary Figure 3B), where the transcription site, marked by NXF1 introns, was positioned near nuclear speckles, while exon labeling emerging from the same transcription site showed a stronger association with the nuclear speckles due to its broader distribution. Additionally, individual transcripts were localized within nuclear speckles (Supplementary Figure 3B). The NXF1 mRNA signal was observed mainly in the nucleoplasm, raising the question of whether NXF1 mRNAs are retained in the nucleus. Our studies showed that NXF1 protein levels increased during infection (Supplementary Figure 4B,C). This supports the infection-induced upregulation of NXF1 transcription (Supplementary Figure 4A). This verifies the formation of active transcription sites and accumulation of mature mRNAs in nuclear speckles, with the mRNA subsequently exported to the cytoplasm for translation. Interestingly, labeling of the NXF1 protein at 3 and 6 hpi revealed that the NXF1 protein is recruited to VRCs (Supplementary Figure 4D,E), probably associating with viral mRNAs en route to export.

**Figure 3:**
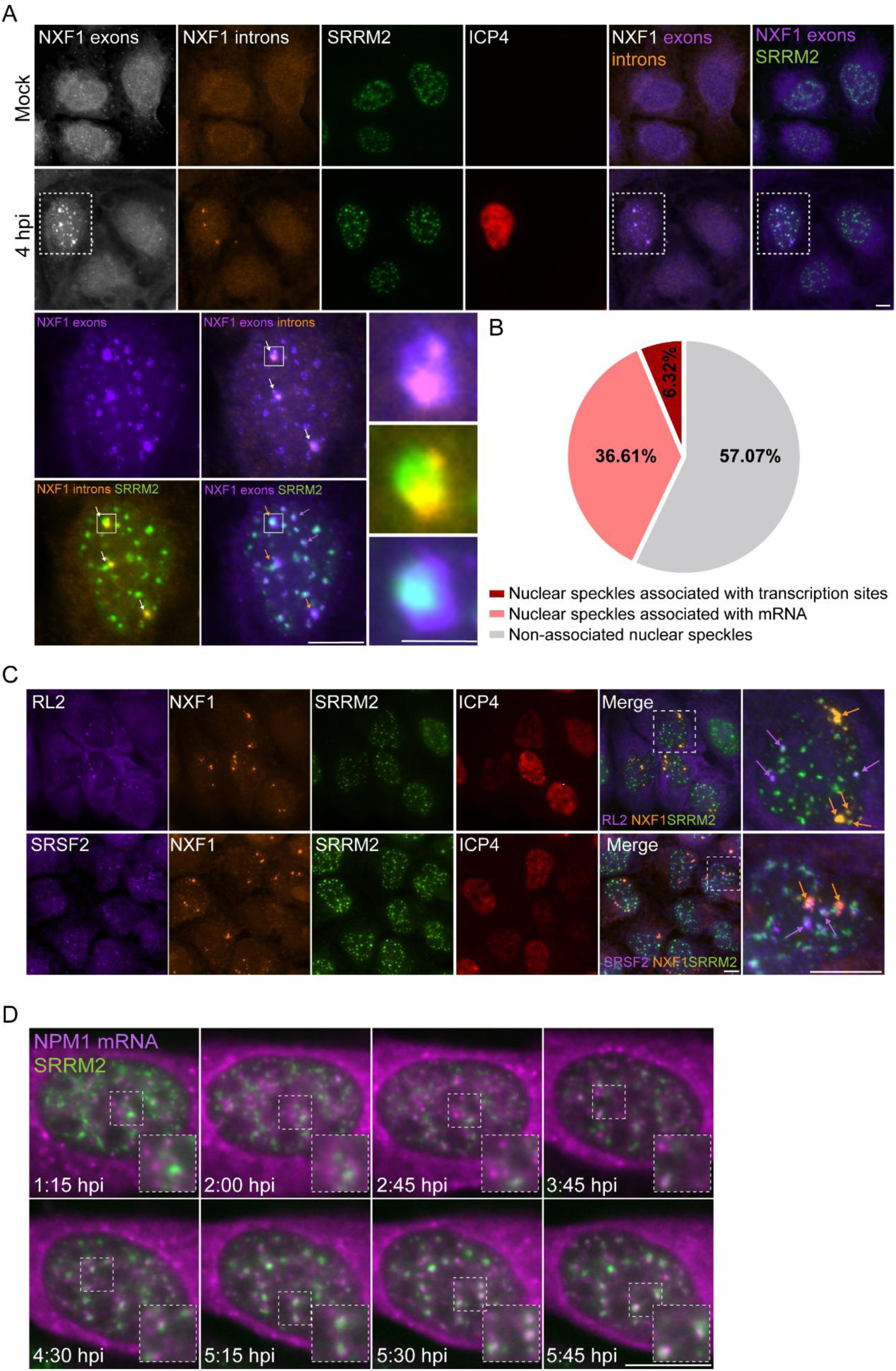
Upregulated host mRNAs are transcribed in close proximity to nuclear speckles and single transcripts localize within them. (A) U2OS cells were mock infected and infected for 4 hrs (MOI=5) and then labeled with fluorescent probes for NXF1 mRNA, using specific probes for NXF1 exons (purple) and introns (orange). SRRM2 marks nuclear speckles (green). ICP4 (red). (B) Quantification of nuclear speckles associated with transcription sites and NXF1 mature transcripts. (C) U2OS cells infected for 4 hrs (MOI=5) and labeled with fluorescent probes for two mRNAs: RS1 (purple) and NXF1 (orange), or SRSF2 (orange) and NXF1 (purple), together with SRRM2 (green) and ICP4 (red). (D) Frames from a time-lapse movie of U2OS cells infected for 6 hrs (MOI=5). Endogenous NPM1 mRNA was tagged using A1-Halo-MCP (purple), and the endogenous SRRM2 protein was fused to a mScarlet fluorescent protein (green). Scale bars= 10 µm.

We examined whether other upregulated cellular genes are associated with nuclear speckles during HSV-1 infection (Supplementary Figure 4A). The genes *SRSF2* encoding the SRSF2 splicing factor, and *NPM1,* encoding the B23 nucleolar protein, were examined. Endogenous SRSF2 mRNAs were detected by smRNA FISH and were also found associated with nuclear speckles during infection (Supplementary Figure 5A). For detecting *NPM1* transcripts, U2OS cells in which the endogenous *NPM1* gene is fused to a Neongreen fluorescent protein and to MS2 sequence repeats by CRISPR knock-in, were used. Cells were transfected with a MS2-coat-protein (MCP) that binds the MS2 sequence repeats and 24 hrs later were infected for 4 hrs. The HaloTag fluorophore (647-halo) was applied to the cells to label the A1-Halo-MCP protein, to detect NPM1 transcripts. The cells were also stained for SRRM2 and ICP4 to observe nuclear speckles and VRCs. NPM1 mRNAs were also found associated with nuclear speckles during HSV-1 infection (Supplementary Figure 5B). Moreover, co-labeling of NXF1 mRNA, SRSF2 mRNA, and nuclear speckles revealed that each actively transcribing gene was associated with specific nuclear speckles and that single transcripts around the site of transcription were associated with nuclear speckles in the proximity of the active gene (Figure 3C). The same phenomenon was seen for the co-labeling of NXF1 and RL2 viral mRNAs (Figure 3C). This shows that active cellular/viral genes are associated with nuclear speckles within their vicinity, and the result of this positioning of genes and nuclear speckles is that transcripts released from the transcription site and diffusing in the nucleoplasm, engage at first with the nuclear speckles nearest to the gene and accumulate there. Later on, as more mRNAs are transcribed, they are found dispersed throughout the nucleus.

To verify the association of host mRNAs with nuclear speckles under viral infection, we performed live-cell imaging to track mRNAs and nuclear speckles in real-time. For live-cell imaging, a similar NPM1-NeonGreen-MS2 cell line was used, in which also the endogenous SRRM2 protein is fused to the mScarlet fluorescent protein (for the detection of nuclear speckles). Cells were transfected with the A1-Halo-MS2-coat protein (MCP), and 24 hrs later infected with HSV-1 ICP4-EYFP and taken for live-cell imaging for over 6 hrs. Early in the infection, NPM1 transcripts were seen diffused in the cytoplasm, however at later stages (4:30 hpi) transcripts were observed associated with nuclear speckles (Figure 3D, Movie S2). In another movie, in which an active transcription site of *NPM1* was observed, we could detect single NPM1 transcripts released from the gene and associating with the nearest nuclear speckles (Movie S3).

### HSV-1 infection induces changes in nuclear speckles abundance, volume, composition and dynamics

MALAT1, a well-characterized long non-coding RNA (lncRNA), is typically enriched in nuclear speckles. We have previously shown that when MALAT1 is lacking in nuclear speckles during transcription inhibition, another lncRNA moves into nuclear speckles (59). Since the nuclear speckles in HSV-1 infected cells showed high accumulation of viral and cellular mRNAs, we examined whether HSV-1 infection affects the localization of MALAT1, as well. Strikingly, infection led to MALAT1 lncRNA translocation from nuclear speckles to the nucleoplasm at 4 hpi (Figure 4A,B). This suggests that the progression of HSV-1 infection disrupts the association of MALAT1 with nuclear speckles.

**Figure 4:**
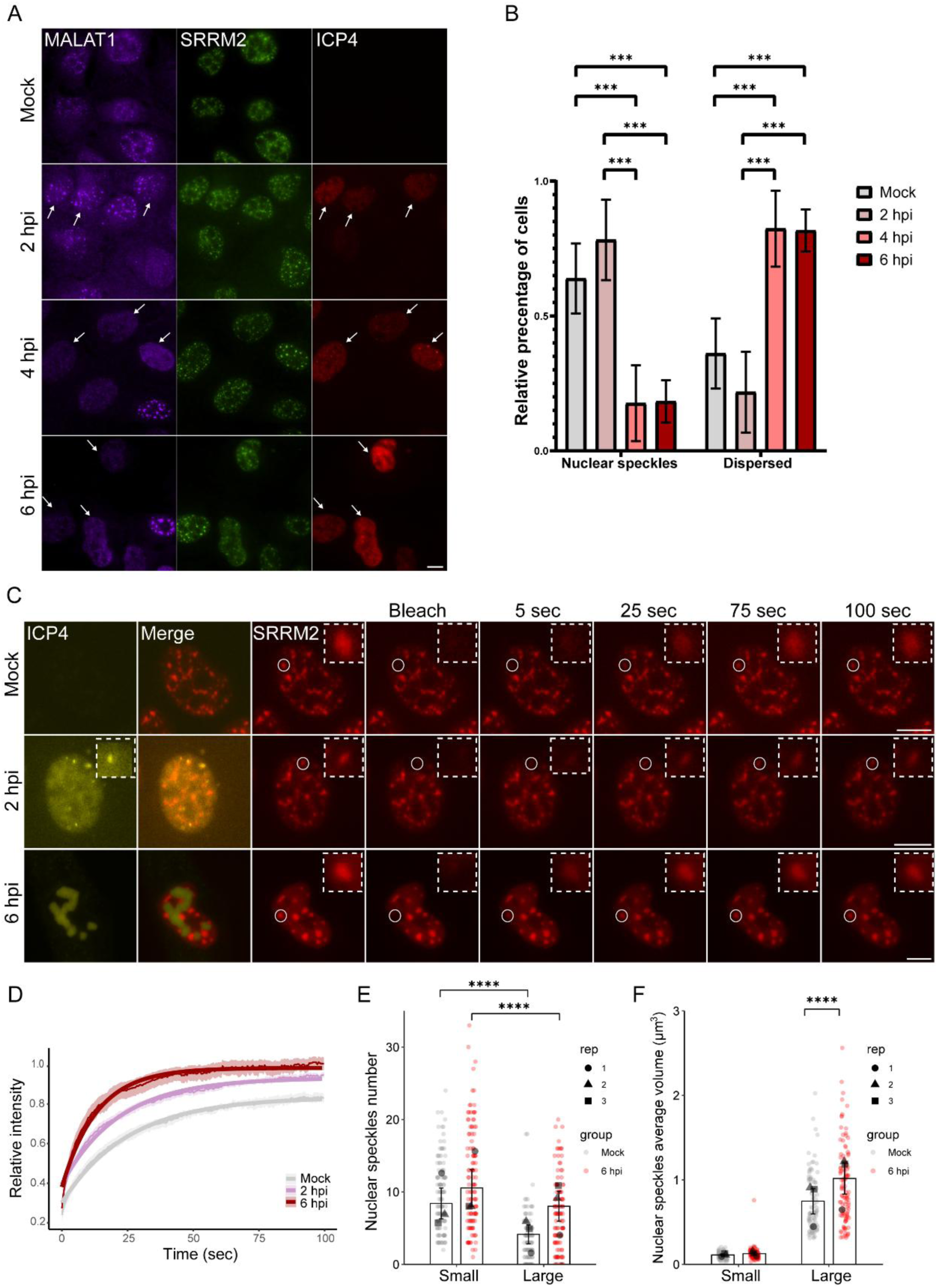
HSV-1 infection induces changes in nuclear speckle composition and dynamics. (A) U2OS cells were infected for 2, 4, and 6 hrs (MOI=5) and lncRNA MALAT1 was labeled by RNA FISH (red). SRRM2 marks nuclear speckles (green). ICP4 protein (red). Scale bar= 10 µm. (B) Quantification of the percentage of cells in which MALAT1 was observed in nuclear speckles versus dispersed in the nucleoplasm is significantly different between mock and 4 hpi. (*P* = 0.0012,), mock and 6 hpi (*P* = 0.0013), 2 hpi and 4 hpi (*P* = 0.0001) and 2 hpi and 6 hpi. (*P* = 0.0001). Control n= 758, 2 hpi n= 1089, 4 hpi n= 1094, 6 hpi n= 1017, in 3 biological replicates. (C) A FRAP experiment on U2OS cells, in which endogenous SRRM2 is fused to mScarlet (red), at: mock infected, 2 hpi and 6 hpi (MOI=5) with HSV-1 ICP4-EYFP (yellow). Scale bar= 10 µm. (D) Plot of FRAP results showing the rate of fluorescence recovery, rate is significantly different between mock and 6 hpi, and 2 hpi and 6 hpi (*P* =0.0206, 0.0248 respectively). The plateau is significantly different between mock and 2 hpi, and mock and 6 hpi (*P* = 0.0372, 0.0112 respectively). Mock n=32, 2 hpi n=39, 6hpi n= 35, in 4 biological replicates. (E) Quantification of nuclear speckle numbers in mock infected cells and 6 hpi cells, categorized to small and large nuclear speckles in each condition (mock vs. 6 hpi). Significant main effects of group (*P* = 5.5e-10) and size (*P* < 2.2e-16); showed no interaction effect (*P* = 0.1926). (F) Quantification of the average volume of nuclear speckles in each cell, categorized to two groups: average volume of the small nuclear speckles (<0.3 µm^3^) and the average volume of the large nuclear speckles (>0.3 µm^3^), with a significant interaction effect (*P* = 2.5e-06), n=120 cells in each condition, in 3 biological replicates (mock and 6 hpi). Post hoc: volume of small nuclear speckles between mock and 6 hpi is not significant (*P*= 0.696), volume of large between mock and 6 hpi is significantly different (*P* = 2.59E-11), volume of small and large nuclear speckles in mock is significantly different (*P*= 1.01e-51), volume of small and large nuclear speckles in 6 hpi is significantly different (*P* = 8.9e-86).

Since the RNA composition of nuclear speckles is modified during HSV-1 infection, we examined whether the dynamics of core nuclear speckles protein is also changing. FRAP analysis of the dynamics of the endogenous SRRM2 protein (fused to the mScarlet fluorescent protein) in nuclear speckles demonstrated that HSV-1 infection significantly affects the dynamics of this nuclear speckle core protein. The nuclear speckles selected for photobleaching were close to ICP4 foci (representing initial viral replication centers) in cells infected for 2 hrs, and to ICP4-associated large VRCs in cells infected for 6 hrs (Figure 4C). The rate constant (m), representing the kinetics of SRRM2 recovery, was significantly higher at 6 hpi compared to both 2 hpi and mock infected cells, indicating faster recovery and increased protein mobility specifically at the later stage of infection (Figure 4D). The plateau values, reflecting the mobile fraction of SRRM2 within the nuclear speckles, were significantly higher at 6 hpi compared to both 2 hpi and mock infected cells. Moreover, the plateau was also significantly higher at 2 hpi compared to mock infected cells, suggesting that HSV-1 infection increases the mobile fraction of SRRM2 in nuclear speckles as early as 2 hpi, with an even greater increase by 6 hpi (Figure 4D). Overall, these results indicate that SRRM2 becomes progressively more mobile and less bound within nuclear speckles during infection. The enhancement in dynamics is observed only at 6 hrs, reflecting distinct time-dependent effects of HSV-1 infection on the dynamics of a nuclear speckle core component in proximity to viral replication centers.

Quantification of nuclear speckle number and volume in mock-infected cells and 6 hpi infected cells demonstrated a significant increase in nuclear speckle abundance under infection conditions (Supplementary Figure 6). Nuclear speckles were classified into two subgroups based on volume: small (<0.3 µm³) and large (>0.3 µm³). Across both conditions, small nuclear speckles were significantly more numerous than large nuclear speckles, with no statistical interaction between infection status and nuclear speckle size distribution (Figure 4E). This means that the increased number of large nuclear speckles observed at 6 hpi reflects the overall increase in nuclear speckle abundance during infection. However, analysis of nuclear speckle volume revealed a significant interaction between treatment (mock vs. 6 hpi) and nuclear speckle size (Figure 4F, and Supplementary Figure 6). Accordingly, at 6 hpi, the mean volume of large speckles was significantly greater than in mock-infected cells. This means that infection has an effect on the large nuclear speckles, which become larger, while small nuclear speckles remain the same in volume.

### Nuclear speckles are crucial hubs for immediate-early viral mRNA processing and export

Close examination of the localization of both viral and cellular mRNA foci associated with nuclear speckles, showed that these foci were predominantly located at the nuclear periphery. This observation prompted us to hypothesize that the accumulation of these transcripts within nuclear speckles could be linked to their subsequent nuclear export. It is known that certain mRNA export factors reside within nuclear speckles, and mRNA may pass through these regions to acquire the necessary export factors (60, 61). To test whether nuclear speckles serve as an intermediate station in the mRNA export pathway, we inhibited nuclear export by overexpressing a dominant-negative form of Dbp5 (Dbp5-DN) (62). The Dbp5 helicase is situated on the cytoplasmic side of the NPC and is required for the release of mRNAs into the cytoplasm. The expression of Dbp5-DN results in an mRNA export block within the NPC (51). Consistent with our hypothesis, we found that HSV-1 infected cells (3 hpi) overexpressing Dbp5-DN exhibited nuclear retention of viral transcripts, particularly within nuclear speckles (Figure 5A,B). This suggests that, prior to export, viral transcripts are likely passing through and processed in nuclear speckles, and when mRNA export is disrupted, the pathway is blocked and the mRNAs remain ‘trapped’ in this compartment.

**Figure 5:**
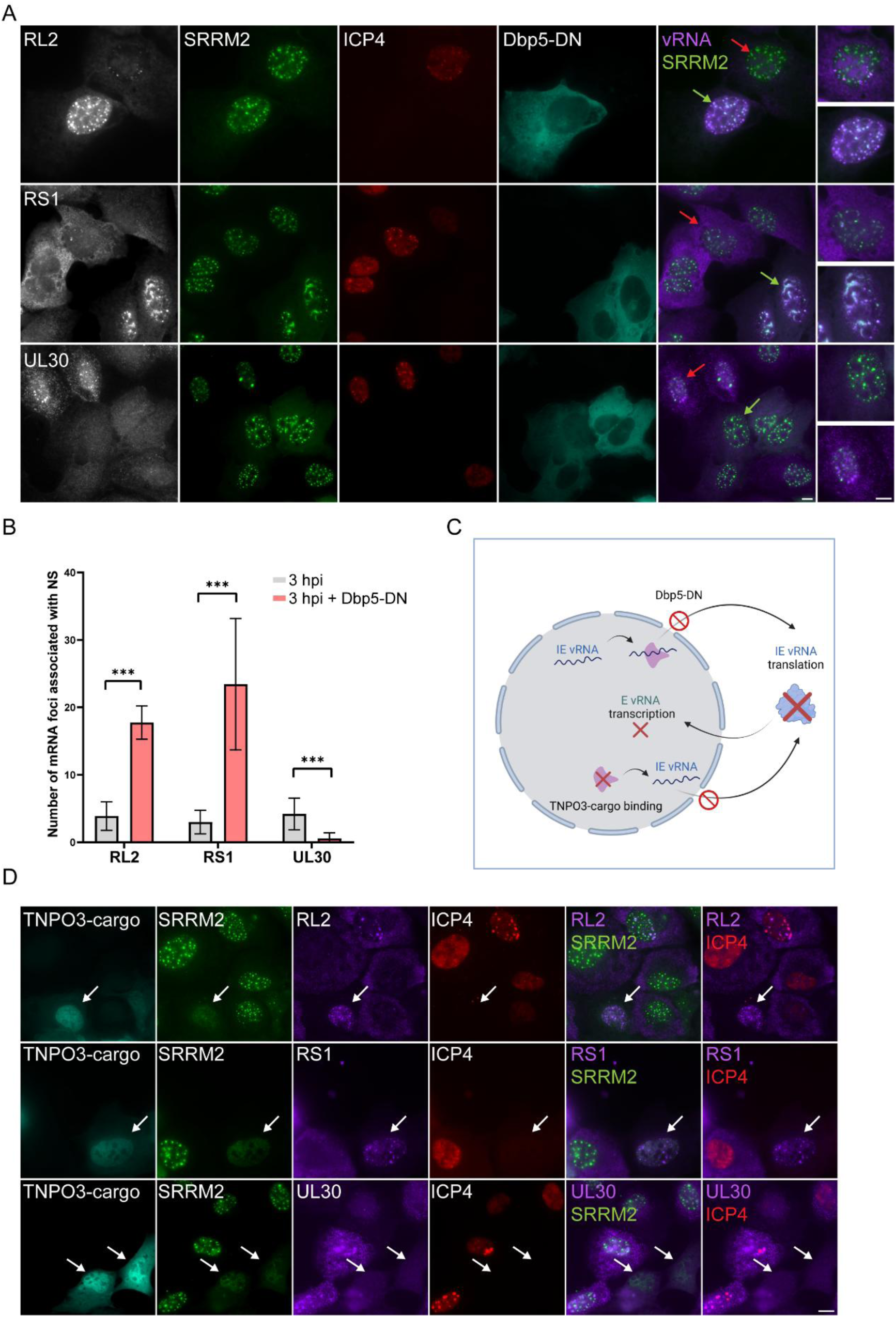
Nuclear speckles are intermediate hubs for IE viral mRNA before export. (A) U2OS cells were transfected with Dbp5-DN-Cerulean (cyan) and 24 hrs later infected for 3 hrs (MOI=5). Viral mRNAs RL2, RS1, and UL30 were labeled (gray) together with the staining of SRRM2 (green) and ICP4 (red). Green arrows refer to Dbp5-DN positive cells, and red arrows refer to Dbp5-DN negative cells. Enlarged images of Dbp5-DN negative/positive cells. Scale bar= 10 µm. (B) Quantification of mRNA accumulation in nuclear speckles at 3 hpi (RL2 n= 131, RS1 n=100, UL30 n= 98) versus 3 hpi + Dbp5-DN (RL2 n= 37, RS1 n= 48, UL30 n=96) is significantly different for RL2 and RS1 (*P* =0.0021, 9.59E-05 respectively). (C) Dbp5-DN blocks IE viral mRNA export and transcripts remain ‘stuck’ in nuclear speckles. Expression of the TNPO3-crago binding domain disperses nuclear speckles and IE viral mRNA are not exported to the cytoplasm. Subsequent translation of IE proteins and E and L viral gene expression is suppressed. (D) U2OS cells were transfected with TNPO3-Cerulean (cyan) and 24 hrs later infected for 3 hrs (MOI=5). Cells were labeled with fluorescent probes for viral mRNA RL2, RS1 and UL30 (gray), together with staining of SRRM2 (green) and ICP4 (red). Scale bar= 10 µm.

As expected, immediate early (IE) mRNAs such as RL2 and RS1 were retained in the nucleus (Figure 5A,B). In contrast, the early (E) gene UL30 mRNA that is expressed after 3-4 hpi (Supplementary Figure 1A, 3A) was barely detectable in Dbp5-DN-expressing cells (Figure 5A), likely because UL30 transcription is dependent on the translation of IE gene products. Since IE mRNAs are not efficiently exported to the cytoplasm in these cells, subsequent transcription and translation of early genes, including UL30, are impaired (Figure 5C). Moreover, co-labeling of the IE transcript RS1 and the E transcript UL30, showed that in Dbp5-DN overexpressing cells RS1 is nuclear retained while UL30 is not detected, compared to Dbp5-DN-negative cells in which RS1 and UL30 were both detected (Supplementary Figure 7A). Interestingly, in rare instances, we did detect UL30 mRNA in cells with Dbp5-DN overexpression, where it was also retained in the nucleus. We speculate that this could occur when Dbp5-DN expression is relatively low or if it is expressed later in the infection cycle, allowing some IE mRNAs to be exported to the cytoplasm to be translated. However, in these cells, UL30 mRNA was not found in nuclear speckles, in contrast to the strong co-localization of RL2 and RS1 mRNAs with these structures (Supplementary Figure 7B). This suggests that nuclear speckles act as intermediate stations for mRNAs on their way to export, specifically for IE transcripts.

We investigated whether UL30 lack of expression in Dbp5-DN cells is due to IE mRNA export block, or due to the export block of host mRNA as well, therefore impairing synthesis of cellular proteins that are needed for viral replication. IE viral transcripts are observed under the mRNA export block, indicating that viral transcription is not impaired. This can be explained by the presence of host proteins which were synthesized prior to Dbp5-DN overexpression, and only new proteins are not translated due to the export block. To further clarify that, we stained the active forms of RNA Pol II (Ser-5 and Ser-2 phosphorylated) in Dbp5-DN overexpressing cells, and indeed found a regular staining pattern, indicating that active transcription is still occurring under this condition (Supplementary Figure 7C). Therefore, we can assume that the lack of expression of UL30 seen in the export blocked cells (Dbp5-DN) is due to the lack of IE proteins caused by the IE mRNA export block, and not due to lack of cellular proteins.

To gain a deeper understanding of the viral mRNA localization within nuclear speckles, we sought to disrupt the nuclear speckle structure and observe its effect on viral mRNA distribution. In a previous study, we demonstrated that overexpression of the cargo-binding domain of TNPO3, a protein involved in the nuclear import of splicing factors (63), leads to the disassembly of nuclear speckles (30). We found that overexpression of the TNPO3 cargo-binding domain caused a dispersion of splicing factors throughout the nucleoplasm. Based on these observations, we hypothesized that if indeed nuclear speckles are important in the pathway of viral mRNA export from the nucleus, then disrupting nuclear speckle structure would cause an export block. Indeed, cells overexpressing this protein and lacking nuclear speckles exhibited retention of the IE viral mRNA RL2 and RS1 in the nucleus at 3 hpi (Figure 5D), indicating that nuclear speckles are required for the export of viral mRNA to the cytoplasm. Consequently, UL30 viral mRNA was not detected in TNPO3-overexpressing cells lacking nuclear speckles (Figure 5C,D). Notably, in these cells with nuclear-retained viral mRNA, the ICP4 viral protein (product of the *RS1* viral gene) was not detected, further supporting the idea that nuclear speckles are involved in the export process of viral transcripts. In conclusion, our findings reveal the path of the IE viral mRNA in the nucleoplasm and the critical role of nuclear speckles in viral mRNA export (Figure 6).

**Figure 6:**
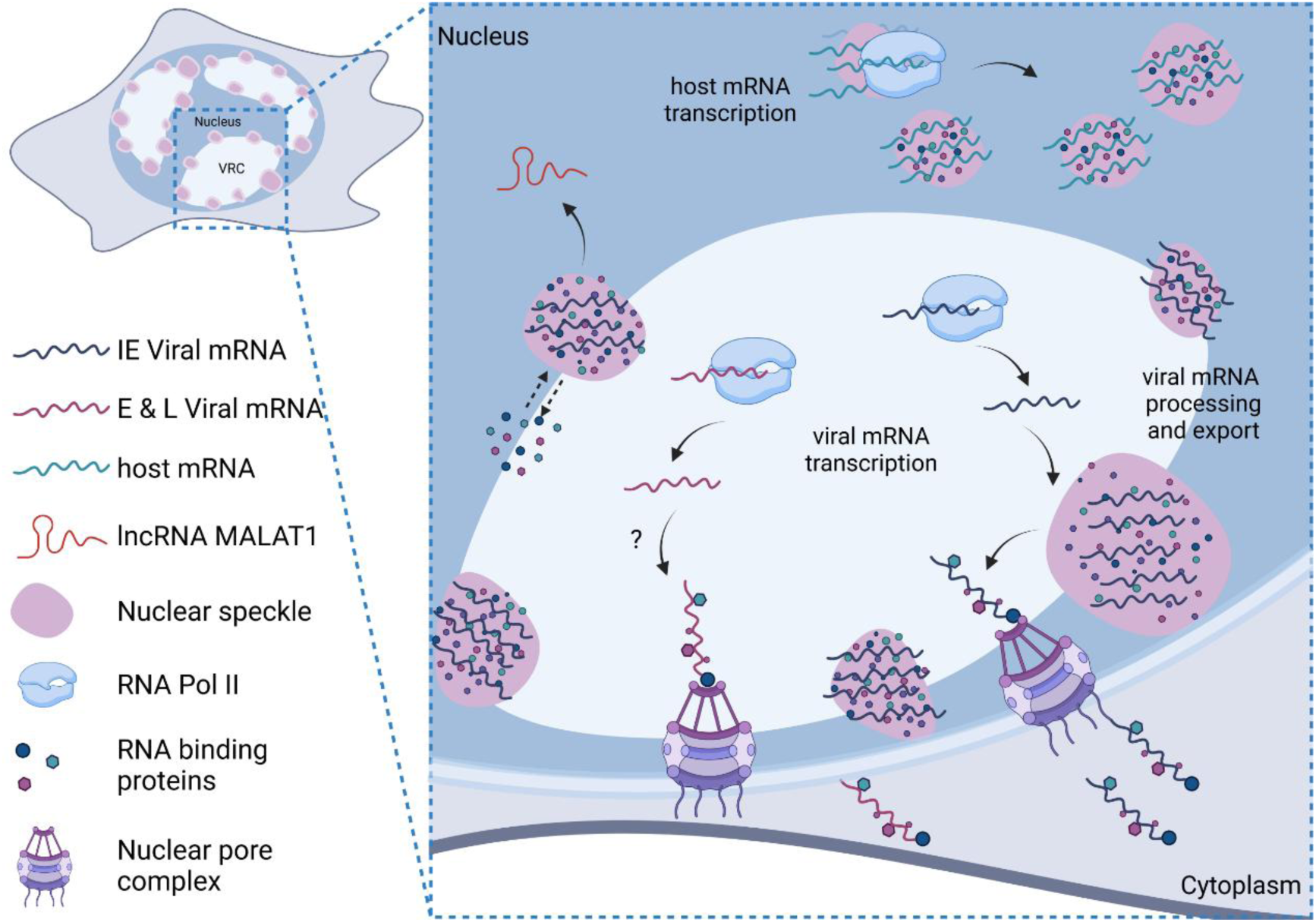
Nuclear speckles and viral and host mRNA during HVS-1 infection. Scheme of viral and host mRNA transcription and their association with nuclear speckles during HSV-1 infection. Viral transcription takes place in the VRCs and IE viral mRNAs traffic to nuclear speckles before export. Host genes transcribe near nuclear speckles during infection and host mRNAs traffic to adjacent nuclear speckles. The lncRNA MALAT1 is redistributed in the nucleoplasm during infection and is not found in nuclear speckles. The dynamics of the nuclear speckles core protein SRRM2 is higher.

## Discussion

In this study, we visualize both viral and cellular gene expression in HSV-1 infected cells and investigate the spatial localization of the expressed transcripts within the nucleus. Our findings demonstrate that IE viral mRNAs, such as RL2 RS1 and US1, accumulate in nuclear speckles after transcription, suggesting a potential role for these structures in viral gene regulation. In contrast, E and L viral mRNAs, such as UL30 and UL36, primarily localize in viral replication compartments, and do not exhibit association with nuclear speckles. This selective accumulation implies that nuclear speckles may function in the post-transcriptional processing of the IE viral genes. Notably, IE viral mRNAs were found in nuclear speckles upon transcription inhibition, confirming that nuclear speckles are not serving as sites of transcription, rather they might be part of the path of viral mRNA trafficking after transcription. Viral transcription inside nuclear speckles would likely require the presence of viral DNA at the VRC rim. The presence of introns of the *RL2* and *US1* viral transcripts in nuclear speckles suggests that post-transcriptional splicing is occurring in this structure. Notably, *RS1* transcripts do not contain introns but still localize to nuclear speckles. This observation is consistent with a study that shows that intronless mRNA passes through nuclear speckles to become export competent (60, 64). Moreover, another nuclear speckle protein, Aly/REF, was previously shown to be involved in the export pathway of intronless viral mRNA (5, 65, 66).

In addition to viral genes, upregulated host mRNAs were also associated with nuclear speckles, further supporting the hypothesis that these nuclear structures play a significant role in gene expression and processing during viral infection, not only for viral genes. This finding aligns with studies showing the positioning of genes near nuclear speckles, highlighting their importance in transcription and splicing enhancement (37, 67–69). Moreover, a recent study on the influenza virus has shown the importance of NS1 viral protein localization to nuclear speckles to inhibit host cell gene expression (70). This further supports the significance of nuclear speckles for gene expression during infection and the strategies viruses apply to make use of them. Like viral genes, which are transcribed in VRCs close to nuclear speckles, the transcription site of host genes is also near nuclear speckles. This underscores the significant role of nuclear speckles in gene expression while demonstrating that even highly active genes are transcribed near, rather than within, nuclear speckles. This observation aligns with previous studies showing a positive correlation between transcription and splicing activity and the proximity of genes to nuclear speckles (34, 71, 72). For example, *HSPA1A* transgenes have been observed to relocate toward nuclear speckles upon heat shock (73), supporting the notion that, under stress conditions, genes move closer to nuclear speckles to enhance their expression, as seen in our study for viral and cellular genes. Moreover, recent findings indicate that nuclear speckles function as central hubs for pre-mRNA 3′-end processing, with over 50% of genes undergoing this process at nuclear speckles (33), reinforcing the idea that transcripts localize to nuclear speckles following transcription. Additionally, while intron sequences of viral mRNA are observed in nuclear speckles, host mRNAs associated with nuclear speckles (not at the site of transcription) are mature transcripts. This means that viral mRNAs may undergo splicing in nuclear speckles, whereas host genes are co-transcriptionally spliced at the site of transcription. Moreover, the examination of two host genes in conjunction with nuclear speckles during infection revealed that each gene accumulates in distinct nuclear speckles. This phenomenon can be explained as the organization of the chromatin during viral infection, where active genes are close to nuclear speckles that contain processing factors, and the mRNAs released from the gene are then trafficked to the nearest nuclear speckles. This is further supported by the observation that mRNA accumulation is found near the site of transcription (identified by introns). Notably, these transcripts are not associated with nuclear speckles under normal conditions. Besides the concentration of viral and host mRNAs in nuclear speckles, HSV-1 induces alterations in the composition and dynamic of these structures. The dispersion of MALAT1 from nuclear speckles and the enhanced mobility of SRRM2 during infection suggest a remodeling of nuclear speckles in response to HSV-1 infection. Notably, higher dynamics of nuclear speckle proteins are not typically attributed to active transcription sites (30). This observation, together with the exclusion of MALAT1 from nuclear speckles and the presence of viral mRNA in these structures at late infection stages (as shown by RNA FISH labeling), suggests that the viral mRNA accumulating in nuclear speckles is not associated with active transcription but rather represents a storage or processing phase. These findings highlight the functional reorganization of nuclear speckles as part of the virus strategy to manipulate cellular resources for its replication and survival. Interestingly, previous studies have shown that the nuclear speckle protein SRSF2, as well as the paraspeckle proteins PSPC1 and p54nrb, along with the scaffold lncRNA NEAT1, undergo nuclear reorganization during HSV-1 infection. These components have been observed to associate with HSV-1 genomic DNA and viral promoters, modulating epigenetic modifications and regulating viral gene expression (5, 65, 66). These findings are consistent with our observations of changes occurring in nuclear speckle composition. Together with our data, showing that viral mRNA transcription occurs in VRCs while some post-transcriptional processing takes place in nuclear speckles, these results suggest that certain nuclear speckle and paraspeckle components may be actively recruited to VRCs to support viral gene expression.

Finally, we show that nuclear speckles may serve as intermediate hubs for the nuclear export of viral mRNAs. Export inhibition using a dominant-negative form of Dbp5, a key mRNA export factor (62), resulted in retention of viral mRNA within nuclear speckles, supporting the notion that these structures serve as intermediate stations in the export process. Expectedly, while IE genes were retained in the nucleus, E and L genes were not expressed. Disruption of the nuclear speckle structures by overexpression of the TNPO3-cargo binding domain (30) led to an export block of the viral IE mRNA, emphasizing the crucial role of nuclear speckles in exporting IE viral mRNAs. Notably, the overexpression of TNPO3-cargo binding domain did not alter the phosphorylation pattern of splicing factors compared to Clk1 overexpression, nor did it prevent their recruitment to active genes or induce a splicing defect, unlike the effects of Pladienolide B (30). Consequently, the disruption of nuclear speckles prevented the expression of E viral genes, as IE mRNAs became trapped in the nucleus and could not induce the transcription of E and L genes. This is further supported by studies proposing nuclear speckles as quality control hubs, ensuring mRNA processing before export (43, 60), which seem to be utilized by viral mRNAs.

While mRNA export block and nuclear speckle disruption demonstrate the significance of nuclear speckles for IE mRNA export, the exact mechanism of viral mRNA export to the cytoplasm remains unraveled. Previous studies have shown the importance of NXF1 in the export of HSV-1 transcripts (74, 75) and influenza transcripts (76). Here, we show that NXF1 is recruited to VRCs and nuclear speckles upon infection, implying its involvement in viral transcript trafficking to nuclear speckles. Additionally, it remains to be determined why only IE mRNAs pass through nuclear speckles before export, while E and L do not. Future studies, including live tracking of viral mRNA, may offer new insights and further our understanding of the spatial organization of gene expression during viral infection.

## Acknowledgments

We are grateful to Ronit Sarid and Inna Kalt (BIU) for kindly providing the Vero cells, their assistance in virus production, and their valuable advice throughout. We also thank Gal Haimovich (Weizmann Institute of Science) for generously sharing the original phage-NLS-HA-tdMCP2-Halo plasmid, and Jennifer I.C. Benichou (BIU) for her assistance with the statistical analysis.

## References

1. A. Calle et al., Nucleolin is required for an efficient herpes simplex virus type 1 infection. J Virol 82, 4762–4773 (2008).

2. R. D. Everett et al., The disruption of ND10 during herpes simplex virus infection correlates with the Vmw110- and proteasome-dependent loss of several PML isoforms. J Virol 72, 6581–6591 (1998).

3. H. Gu, B. Roizman, The degradation of promyelocytic leukemia and Sp100 proteins by herpes simplex virus 1 is mediated by the ubiquitin-conjugating enzyme UbcH5a. Proc. Natl. Acad. Sci. U S A 100, 8963–8968 (2003).

4. M. K. Chelbi-Alix, H. de The, Herpes virus induced proteasome-dependent degradation of the nuclear bodies-associated PML and Sp100 proteins. Oncogene 18, 935–941 (1999).

5. K. Li, Z. Wang, Speckles and paraspeckles coordinate to regulate HSV-1 genes transcription. Commun Biol 4, 1207 (2021).

6. R. M. Sandri-Goldin, M. K. Hibbard, M. A. Hardwicke, The C-terminal repressor region of herpes simplex virus type 1 ICP27 is required for the redistribution of small nuclear ribonucleoprotein particles and splicing factor SC35; however, these alterations are not sufficient to inhibit host cell splicing. J Virol 69, 6063–6076 (1995).

7. C. J. Lukonis, S. K. Weller, Formation of herpes simplex virus type 1 replication compartments by transfection: requirements and localization to nuclear domain 10. J Virol 71, 2390–2399 (1997).

8. T. E. Martin, S. C. Barghusen, G. P. Leser, P. G. Spear, Redistribution of nuclear ribonucleoprotein antigens during herpes simplex virus infection. J Cell Biol 105, 2069–2082 (1987).

9. M. Simpson-Holley, R. C. Colgrove, G. Nalepa, J. W. Harper, D. M. Knipe, Identification and functional evaluation of cellular and viral factors involved in the alteration of nuclear architecture during herpes simplex virus 1 infection. J Virol 79, 12840–12851 (2005).

10. D. T. McSwiggen et al., Evidence for DNA-mediated nuclear compartmentalization distinct from phase separation. eLife 8 (2019).

11. E. Caragliano, W. Brune, J. B. Bosse, Herpesvirus Replication Compartments: Dynamic Biomolecular Condensates? Viruses 14 (2022).

12. L. Chang et al., Herpesviral replication compartments move and coalesce at nuclear speckles to enhance export of viral late mRNA. Proc. Natl. Acad. Sci. U S A 108, E136–144 (2011).

13. E. Tomer et al., Coalescing replication compartments provide the opportunity for recombination between coinfecting herpesviruses. FASEB J 33, 9388–9403 (2019).

14. H. Xu et al., Two-Color CRISPR Imaging Reveals Dynamics of Herpes Simplex Virus 1 Replication Compartments and Virus-Host Interactions. J Virol 96, e0092022 (2022).

15. V. Aho et al., Chromatin organization regulates viral egress dynamics. Scientific reports 7, 3692 (2017).

16. K. Monier, J. C. Armas, S. Etteldorf, P. Ghazal, K. F. Sullivan, Annexation of the interchromosomal space during viral infection. Nat. Cell Biol. 2, 661–665 (2000).

17. M. Simpson-Holley, J. Baines, R. Roller, D. M. Knipe, Herpes simplex virus 1 U(L)31 and U(L)34 gene products promote the late maturation of viral replication compartments to the nuclear periphery. J Virol 78, 5591–5600 (2004).

18. T. Chew, K. E. Taylor, K. L. Mossman, Innate and adaptive immune responses to herpes simplex virus. Viruses 1, 979–1002 (2009).

19. A. Esclatine, B. Taddeo, L. Evans, B. Roizman, The herpes simplex virus 1 UL41 gene-dependent destabilization of cellular RNAs is selective and may be sequence-specific. Proc. Natl. Acad. Sci. U S A 101, 3603–3608 (2004).

20. A. J. Guise, H. G. Budayeva, B. A. Diner, I. M. Cristea, Histone deacetylases in herpesvirus replication and virus-stimulated host defense. Viruses 5, 1607–1632 (2013).

21. K. L. Mossman et al., Herpes simplex virus triggers and then disarms a host antiviral response. J Virol 75, 750–758 (2001).

22. K. Nystrom et al., Real time PCR for monitoring regulation of host gene expression in herpes simplex virus type 1-infected human diploid cells. Journal of virological methods 118, 83–94 (2004).

23. N. Van Opdenbosch, H. Favoreel, G. R. Van de Walle, Histone modifications in herpesvirus infections. Biology of the cell / under the auspices of the European Cell Biology Organization 104, 139–164 (2012).

24. J. Xiang, C. Fan, H. Dong, Y. Ma, P. Xu, A CRISPR-based rapid DNA repositioning strategy and the early intranuclear life of HSV-1. eLife 12 (2023).

25. D. L. Spector, A. I. Lamond, Nuclear speckles. Cold Spring Harb Perspect Biol 3 (2011).

26. L. Galganski, M. O. Urbanek, W. J. Krzyzosiak, Nuclear speckles: molecular organization, biological function and role in disease. Nucleic Acids Res 45, 10350–10368 (2017).

27. G. P. Faber, S. Nadav-Eliyahu, Y. Shav-Tal, Nuclear speckles - a driving force in gene expression. J. Cell Sci. 135 (2022).

28. P. Chaturvedi, A. S. Belmont, Nuclear speckle biology: At the cross-roads of discovery and functional analysis. Curr Opin Cell Biol 91, 102438 (2024).

29. S. Huang, D. L. Spector, Will the real splicing sites please light up? Curr. Biol. 2, 188–190 (1992).

30. H. Hochberg-Laufer et al., Availability of splicing factors in the nucleoplasm can regulate the release of mRNA from the gene after transcription. PLoS genetics 15, e1008459 (2019).

31. T. Misteli et al., Serine phosphorylation of SR proteins is required for their recruitment to sites of transcription in vivo. J Cell Biol 143, 297–307 (1998).

32. T. D. Williams et al., mRNA export factors store nascent transcripts within nuclear speckles as an adaptive response to transient global inhibition of transcription. Mol. Cell 85, 117–131 e117 (2025).

33. Y. Yoon et al., RBBP6 anchors pre-mRNA 3’ end processing to nuclear speckles for efficient gene expression. Mol. Cell 85, 555–570 e558 (2025).

34. Y. Chen et al., Mapping 3D genome organization relative to nuclear compartments using TSA-Seq as a cytological ruler. J Cell Biol 10.1083/jcb.201807108 (2018).

35. A. S. Belmont, Nuclear Compartments: An Incomplete Primer to Nuclear Compartments, Bodies, and Genome Organization Relative to Nuclear Architecture. Cold Spring Harb Perspect Biol 10.1101/cshperspect.a041268 (2021).

36. Y. Brody et al., The in vivo kinetics of RNA polymerase II elongation during co-transcriptional splicing. PLoS Biol 9, e1000573 (2011).

37. P. Bhat et al., Genome organization around nuclear speckles drives mRNA splicing efficiency. Nature 629, 1165–1173 (2024).

38. S. Hu, P. Lv, Z. Yan, B. Wen, Disruption of nuclear speckles reduces chromatin interactions in active compartments. Epigenetics Chromatin 12, 43 (2019).

39. L. Escudero-Paunetto, L. Li, F. P. Hernandez, R. M. Sandri-Goldin, SR proteins SRp20 and 9G8 contribute to efficient export of herpes simplex virus 1 mRNAs. Virology 401, 155–164 (2010).

40. M. S. Friedl et al., HSV-1 and influenza infection induce linear and circular splicing of the long NEAT1 isoform. PLoS One 17, e0276467 (2022).

41. K. L. Harper et al., Virus-modified paraspeckle-like condensates are hubs for viral RNA processing and their formation drives genomic instability. Nature communications 15, 10240 (2024).

42. E. Alkalay et al., The Sub-Nuclear Localization of RNA-Binding Proteins in KSHV-Infected Cells. Cells 9 (2020).

43. A. Mor et al., Influenza virus mRNA trafficking through host nuclear speckles. Nat Microbiol 1, 16069 (2016).

44. A. C. Francis et al., HIV-1 replication complexes accumulate in nuclear speckles and integrate into speckle-associated genomic domains. Nature communications 11, 3505 (2020).

45. P. Bhat et al., Influenza virus mRNAs encode determinants for nuclear export via the cellular TREX-2 complex. Nature communications 14, 2304 (2023).

46. J. Fueller et al., CRISPR-Cas12a-assisted PCR tagging of mammalian genes. J Cell Biol 219 (2020).

47. M. Sandbaumhuter et al., Cytosolic herpes simplex virus capsids not only require binding inner tegument protein pUL36 but also pUL37 for active transport prior to secondary envelopment. Cell Microbiol 15, 248–269 (2013).

48. L. Ivanova et al., Conserved Tryptophan Motifs in the Large Tegument Protein pUL36 Are Required for Efficient Secondary Envelopment of Herpes Simplex Virus Capsids. J Virol 90, 5368–5383 (2016).

49. A. Buch et al., Inner tegument proteins of Herpes Simplex Virus are sufficient for intracellular capsid motility in neurons but not for axonal targeting. PLoS pathogens 13, e1006813 (2017).

50. A. Mor et al., Dynamics of single mRNP nucleocytoplasmic transport and export through the nuclear pore in living cells. Nat. Cell Biol. 12, 543–552 (2010).

51. R. Ben-Yishay et al., Imaging within single NPCs reveals NXF1’s role in mRNA export on the cytoplasmic side of the pore. J Cell Biol 218, 2962–2981 (2019).

52. N. Tsanov et al., smiFISH and FISH-quant - a flexible single RNA detection approach with super-resolution capability. Nucleic Acids Res 44, e165 (2016).

53. N. Otsu, A Threshold Selection Method from Gray-Level Histograms. *IEEE Transactions on Systems*, Man, and Cybernetics 9, 62–66 (1979).

54. J. N. Kapur, P. K. Sahoo, A. K. C. Wong, A new method for gray-level picture thresholding using the entropy of the histogram. Computer Vision, Graphics, and Image Processing 29, 273–285 (1985).

55. S. Leclerc et al., Progression of herpesvirus infection remodels mitochondrial organization and metabolism. PLoS pathogens 20, e1011829 (2024).

56. D. Tombacz et al., Multiple Long-Read Sequencing Survey of Herpes Simplex Virus Dynamic Transcriptome. Front Genet 10, 834 (2019).

57. C. C. Friedel et al., Dissecting Herpes Simplex Virus 1-Induced Host Shutoff at the RNA Level. J Virol 95 (2021).

58. H. Shenasa, D. L. Bentley, Pre-mRNA splicing and its cotranscriptional connections. Trends in genetics: TIG 39, 672–685 (2023).

59. S. E. Hasenson et al., The Association of MEG3 lncRNA with Nuclear Speckles in Living Cells. Cells 11 (2022).

60. K. Wang et al., Intronless mRNAs transit through nuclear speckles to gain export competence. J Cell Biol 10.1083/jcb.201801184 (2018).

61. S. Gao et al., Nuclear speckle integrity and function require TAO2 kinase. Proc. Natl. Acad. Sci. U S A 119, e2206046119 (2022).

62. C. A. Hodge et al., The Dbp5 cycle at the nuclear pore complex during mRNA export I: dbp5 mutants with defects in RNA binding and ATP hydrolysis define key steps for Nup159 and Gle1. Genes Dev 25, 1052–1064 (2011).

63. G. N. Maertens et al., Structural basis for nuclear import of splicing factors by human Transportin 3. Proc. Natl. Acad. Sci. U S A 111, 2728–2733 (2014).

64. I. H. Chen, K. S. Sciabica, R. M. Sandri-Goldin, ICP27 interacts with the RNA export factor Aly/REF to direct herpes simplex virus type 1 intronless mRNAs to the TAP export pathway. J Virol 76, 12877–12889 (2002).

65. Z. Wang et al., Serine/Arginine-rich Splicing Factor 2 Modulates Herpes Simplex Virus Type 1 Replication via Regulating Viral Gene Transcriptional Activity and Pre-mRNA Splicing. J Biol Chem 291, 26377–26387 (2016).

66. Z. Wang et al., NEAT1 modulates herpes simplex virus-1 replication by regulating viral gene transcription. Cell Mol Life Sci 74, 1117–1131 (2017).

67. J. Kim, N. C. Venkata, G. A. Hernandez Gonzalez, N. Khanna, A. S. Belmont, Gene expression amplification by nuclear speckle association. J Cell Biol 219 (2020).

68. J. Wu et al., Dynamics of RNA localization to nuclear speckles are connected to splicing efficiency. Sci Adv 10, eadp7727 (2024).

69. K. A. Alexander et al., p53 mediates target gene association with nuclear speckles for amplified RNA expression. Mol. Cell 81, 1666–1681 e1666 (2021).

70. W. Nacken, J. Mayr, A. Schreiber, S. Ludwig, Influenza A virus NS1 suppresses nuclear speckles promoted gene expression by inhibition of transcription. Npj Viruses 3, 46 (2025).

71. Y. Chen, A. S. Belmont, Genome organization around nuclear speckles. Curr Opin Genet Dev 55, 91–99 (2019).

72. F. Ding, M. B. Elowitz, Constitutive splicing and economies of scale in gene expression. Nat Struct Mol Biol 26, 424–432 (2019).

73. N. Khanna, Y. Hu, A. S. Belmont, HSP70 transgene directed motion to nuclear speckles facilitates heat shock activation. Curr. Biol. 24, 1138–1144 (2014).

74. L. A. Johnson, L. Li, R. M. Sandri-Goldin, The cellular RNA export receptor TAP/NXF1 is required for ICP27-mediated export of herpes simplex virus 1 RNA, but the TREX complex adaptor protein Aly/REF appears to be dispensable. J Virol 83, 6335–6346 (2009).

75. L. A. Johnson, R. M. Sandri-Goldin, Efficient nuclear export of herpes simplex virus 1 transcripts requires both RNA binding by ICP27 and ICP27 interaction with TAP/NXF1. J Virol 83, 1184–1192 (2009).

76. K. Zhang et al., Cellular NS1-BP protein interacts with the mRNA export receptor NXF1 to mediate nuclear export of influenza virus M mRNAs. J Biol Chem 300, 107871 (2024).

